# NeuroRA: A Python Toolbox of Representational Analysis from Multi-modal Neural Data

**DOI:** 10.1101/2020.03.25.008086

**Authors:** Zitong Lu, Yixuan Ku

**Affiliations:** Guangdong Provincial Key Laboratory of Social Cognitive Neuroscience and Mental Health, Department of Psychology, Sun Yat-sen University, Guangzhou, China; Peng Cheng Laboratory, Shenzhen, China; Shanghai Key Laboratory of Brain Functional Genomics, Shanghai Changning-ECNU Mental Health Center, School of Psychology and Cognitive Science, East China Normal University, Shanghai, 200062, China; NYU-ECNU Institute of Brain and Cognitive Science, NYU Shanghai and Collaborative Innovation Center for Brain Science, Shanghai, China

**Author notes:** correspondence: Yixuan Ku, ResearcherID: D-4063-2018.

**Keywords:** Representational similarity analysis, multivariate pattern analysis, multi-modal, python, correlation analysis

## Abstract

In studies of cognitive neuroscience, multivariate pattern analysis (MVPA) is widely used as it offers richer information than traditional univariate analysis. Representational similarity analysis (RSA), as one method of MVPA, has become an effective decoding method based on neural data by calculating the similarity between different representations in the brain under different conditions. Moreover, RSA is suitable for researchers to compare data from different modalities, and even bridge data from different species. However, previous toolboxes have been made to fit for specific datasets. Here, we develop a novel and easy-to-use toolbox based on Python named NeuroRA for representational analysis. Our toolbox aims at conducting cross-modal data analysis from multi-modal neural data (e.g. EEG, MEG, fNIRS, ECoG, sEEG, neuroelectrophysiology, fMRI), behavioral data, and computer simulated data. Compared with previous software packages, our toolbox is more comprehensive and powerful. By using NeuroRA, users can not only calculate the representational dissimilarity matrix (RDM), which reflects the representational similarity between different conditions, but also conduct a representational analysis among different RDMs to achieve a cross-modal comparison. In addition, users can calculate neural pattern similarity, spatiotemporal pattern similarity (STPS) and inter-subject correlation (ISC) with this toolbox. NeuroRA also provides users with functions performing statistical analysis, storage and visualization of results. We introduce the structure, modules, features, and algorithms of NeuroRA in this paper, as well as examples applying the toolbox in published datasets.

## Introduction

In recent years, research on brain science based on neural data has shifted from univariate analysis towards multivariate pattern analysis (MVPA) (Norman et al., 2006). In contrast to the former, the latter accounts for the population coding for neurons. The decoding of neural activity can help scientists better understand the encoding process of neurons. As in David Marr’s model, representation bridges the gap between a computation goal and implementation machinery (Marr, 1982). Representational similarity analysis (RSA) (Kriegeskorte et al., 2008a) is an effective MVPA method that can successfully describe the relationship between representations of different modalities of data, bridging gaps between human and animals. Therefore, RSA has been rapidly applied in investigating various cognitive functions, including perception (Evans et al., 2015; Henriksson et al., 2019), memory (Xue et al., 2010), language (Chen et al., 2016), and decision-making (Yan et al., 2016).

With the technological development in brain science, various neural recording methods have emerged rapidly. Noninvasive neurophysiological recordings such as electroencephalography (EEG) and magnetoencephalography (MEG) with high temporal resolution, and neuroimaging methods such as functional near-infrared spectroscopy (fNIRS) and functional magnetic resonance imaging (fMRI) with high spatial resolution, have been widely used for basic research. Meanwhile, invasive techniques such as electrocorticography (ECoG), stereo-electro-encephalography (sEEG), and neuroelectrophysiology have been applied to patients or non-human primates. The interpretation of results across different recording modalities becomes difficult. The RSA method, however, uses a representation dissimilarity matrix (RDM) to bridge data from different modalities. For example, studies have attempted to combine fMRI results with electrophysiological results (Kriegeskorte et al., 2008b) or MEG results with electrophysiological results (Cichy et al., 2014). Moreover, it can connect behavioral and neural representational matrices (Wang et al., 2018). Furthermore, with the rapid development of artificial intelligence (AI) (Jordan and Mitchell, 2015; Kriegeskorte and Golan, 2019), RSA can be used to compare representations in artificial neural networks (ANN) with those in EEG (Greene and Hansen, 2018). In summary, RSA is a useful tool to combine the results of behavior and multi-modal neural data, which can lead to a better understanding of the brain, and even further, can help us establish a clearer link between the brain and artificial intelligence (Khaligh-Razavi and Kriegeskorte, 2014; Güçlü and van Gerven, 2015; Eickenberg et al., 2017; Greene and Hansen, 2018b; Kuzovkin et al., 2018).

Some existing tools for RSA include a module in PyMVPA (Hanke et al., 2009), a toolbox for RSA by Kriegeskorte (Nili et al., 2014) and an example in MNE-Python (Gramfort et al., 2013). However, they all have some shortcomings. MNE can only perform RSA for MEG and EEG data in one example. PyMVPA can only implement some basic functions, such as calculating the correlation coefficient and data conversion. Kriegeskorte’s toolbox attached to their paper is designed mainly based on fMRI data and users need to be proficient in MATLAB (Kriegeskorte et al., 2008b), which makes it difficult to generate to other datasets. We considered build a comprehensive and universal toolbox for RSA, and Python was chosen as a suitable programming language. Python is a rapidly rising programming language having great advantages for scientific computing (Sanner, 1999; Koepke, 2011). Because of its strong expansibility, it is more accommodating to use Python for computing and incorporate a toolbox inside it. NumPy (Van et al., 2011), Scikit-learn (Pedregosa et al., 2013), and some other extensions can realize and simplify various basic computing functions. Thus, a number of researchers select Python to develop toolkits in psychology and neuroscience, such as PsychoPy (Peirce, 2007) for designing psychological experiment programs, MNE-Python for EEG/MEG data analysis, and PyMVPA for utilizing MVPA in data from different modalities.

In the present toolbox, we have developed a novel and easy-to-use Python toolbox, NeuroRA (neural representational analysis), for comprehensive representation analysis. NeuroRA aims at using powerful computational resources with Python and conducting cross-modal data analyses for various types of neural data (e.g. EEG, MEG, fNIRS, fMRI), as well as behavioral data and computer stimulation data. In addition to traditional functions of RSA, NeuroRA also includes some specialized functions of representational analysis in published papers across several laboratories, such as neural pattern similarity (NPS), spatiotemporal pattern similarity (STPS) (Xue et al., 2010; Lu et al., 2015) and inter-subject correlation (ISC) (Hasson et al., 2004). NeuroRA requires several basic Python packages to function, including NumPy, SciPy, Matplotlib (Hunter, 2007), Nibabel (Brett et al., 2016), Nilearn and MNE-Python. In the following sections, we detail the structure and function of NeuroRA and further apply it to the open dataset of a MEG study (Cichy et al., 2014) and a fMRI study (Haxby 2001) to guide users to apply NeuroRA.

## Overview of NeuroRA

The structure and functions of NeuroRA are illustrated in **Figure 1**. It can analyze all types of neural (including EEG, MEG, fNIRS, ECoG, sEEG, electrophysiological and fMRI data) and behavioral data. By utilizing the powerful computational toolbox in Python, NeuroRA gives users the ability to mine neural data thoroughly and efficiently.

**Figure 1.**
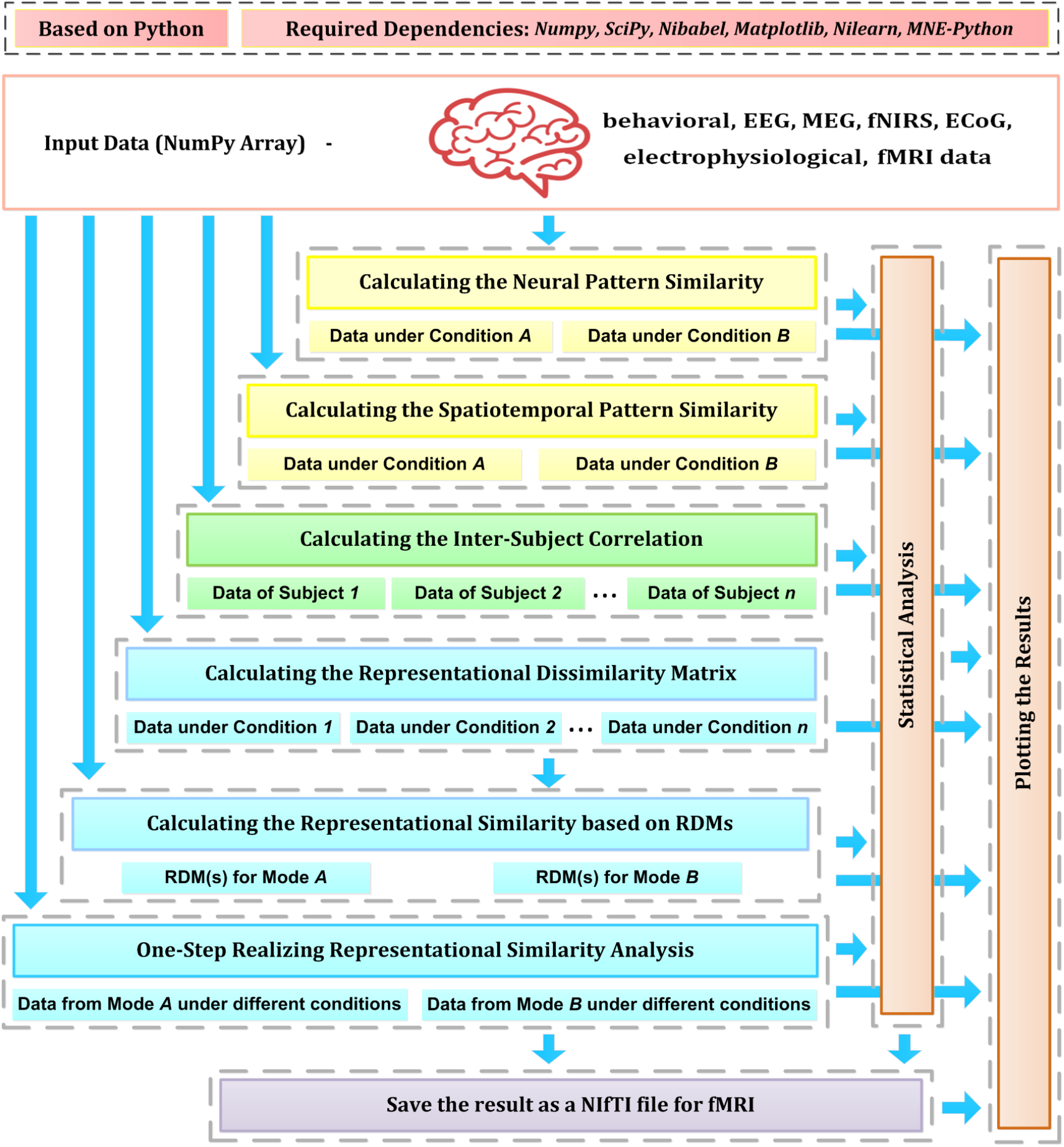
Overview of NeuroRA. NeuroRA is a Python-based toolbox and requires some extension packages, including NumPy, SciPy, Matplotlib, Nilearn and MNE-Python. It contains several main functions: calculating neural pattern similarity (NPS), spatiotemporal pattern similarity (STPS), inter-subject correlation (ISC), and representation dissimilarity matrix (RDM), comparing representations among different modalities using RDMs, statistical analysis, saving results as a NIfTI file for fMRI data, and plotting the results. The blue arrows indicate the data flow. The specific implementation of these features is listed in the main text.

NeuroRA provides abundant functions. First, NPS function reflects the correlation of brain activities induced under two different conditions. Second, STPS function reflects the representational similarity across different space and time points. Third, ISC function reflects the similarity of brain activity among multiple subjects under the same condition. Fourth, RDM function reflects the representation similarity between different conditions/stimuli with neural data from a given modality. All values in the matrix are normalized, and the value at any point in the matrix reflects the dissimilarity of the data representation under the two conditions corresponding to the row and column respectively. Points on the diagonal use the same data under the same conditions, thus the dissimilarity value is 0. Fifth, NeuroRA performs a correlation analysis between RDMs from different modalities to compare representations across modalities. This procedure can be applied according to different parameters; for example, the calculation can be applied for each subject, for each channel, for each time-point, or a combination of all of them.

In addition to calculating the above values, NeuroRA provides a statistical module to perform statistical analysis based on those values and a visualization module to plot the results, such as RDMs, representational similarities over time, and RSA-results for fMRI. Also, NeuroRA provides a unique approach to save the result of representational analysis back to fMRI widely used format, i.e. a NIfTI file obtained with user defined output-threshold.

The pre-required packages for NeuroRA include NumPy, SciPy, Matplotlib, Nilearn and MNE-Python, which are checked and automatically downloaded by installing NeuroRA. NumPy assists with matrix-based computation. SciPy helps with basic statistical analysis. Matplotlib is employed for the plotting functions. NiBabel is used to read and generate NIfTI files. Users can download NeuroRA through only one line of command: *pip install neurora*. The website for our toolbox is https://neurora.github.io/NeuroRA/, and the GitHub URL for its source code is https://github.com/neuora/NeuroRA.

## Data Structures in NeuroRA

The calculations in NeuroRA are all based on multidimensional matrices, including deformation, transposition, decomposition, standardization, addition, and subtraction. The data type in NeuroRA is *ndarray*, an N-dimensional array class of NumPy. Therefore, users first convert their neural data into a matrix (*ndarray* type) as the input of NeuroRA, with information on the different dimensions of the matrix, such as the number of subjects, number of conditions, number of channels, and size of the image (see instructions in the software for details). Here, we give users some feasible methods for data conversion for different kinds of neural data in **Tabel 1**. The outputs of the functions in NeuroRA are square matrices with the same dimensions as the input matrix. The input and output data structures of the NeuroRA functions are shown in the tutorial attached in the website.

**Table 1.**
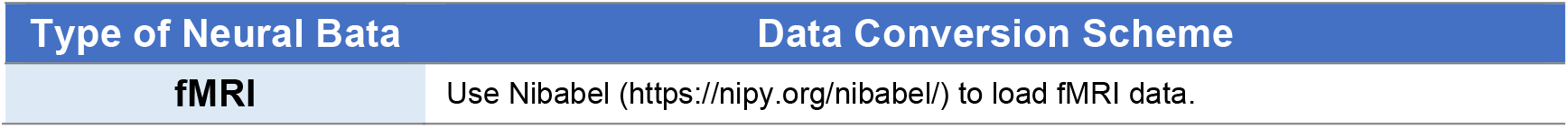
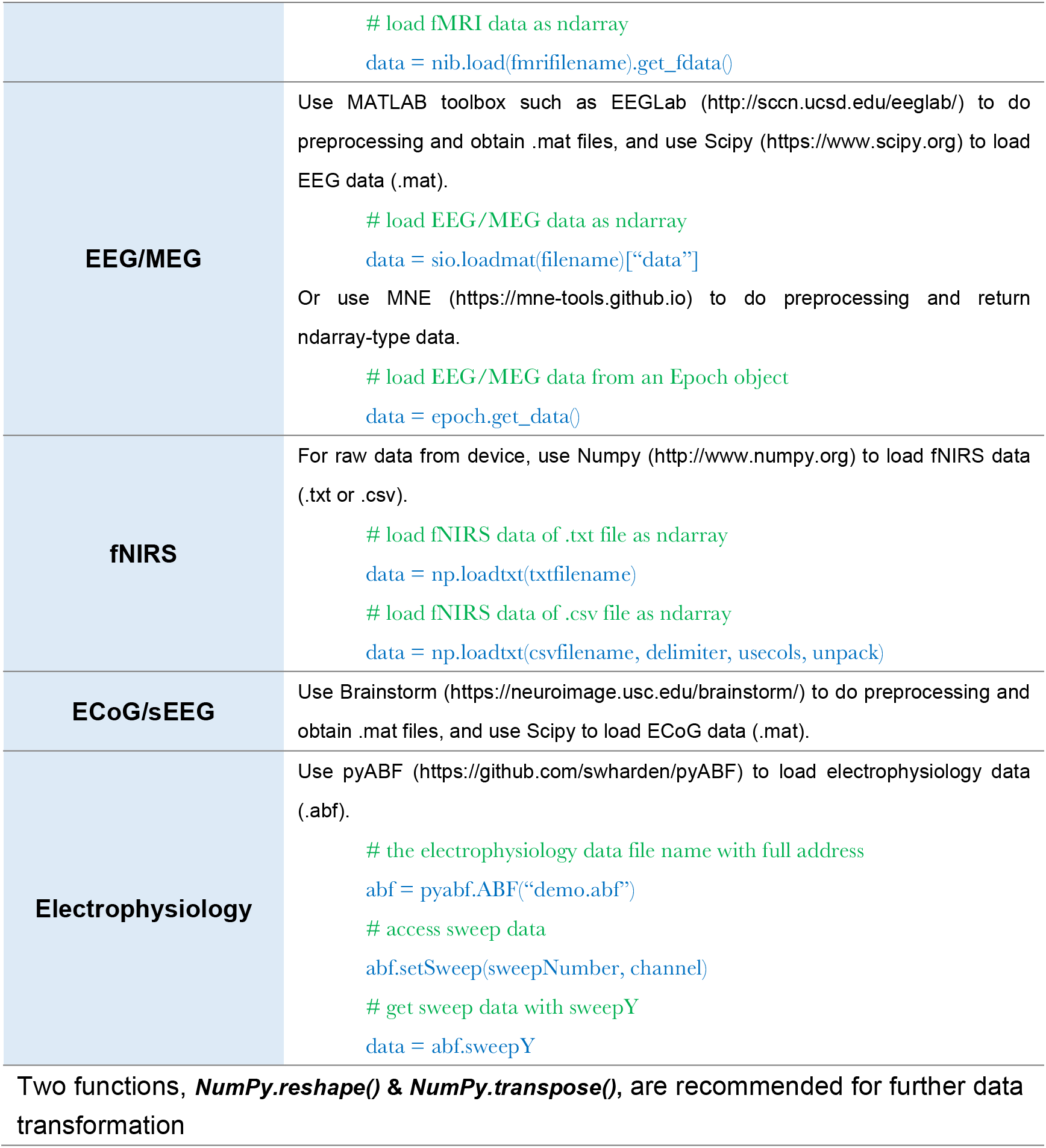
Recommeded data conversion scheme

### NeuroRA’s Modules and Features

NeuroRA attains various functions to process the representational analysis. Usually, data must be processed in multi-step ways, and this toolkit highly integrates these intermediate processes, making it easy to implement. In NeuroRA, only a simple function is required to complete the following processes. Users can obtain the required results after a necessary conversion of the data format.

Meanwhile, we attempt to add some adjustable parameters to meet the calculation requirements for different experiments and different modalities of data. Users can flexibly change the input parameters in the function to match their data format and computing goals.

NeuroRA mainly includes the following core modules, and more modules could be added in the future or as requested.

*nps_cal*: A module to calculate the neural pattern similarity based on neural data.
*stps_cal*: A module to calculate the spatiotemporal pattern similarity based on neural data.
*isc_cal*: A module to calculate the inter-subject correlation based on neural data.
*rdm_cal*: A module to calculate RDM based on multi-modal neural data.
*rdm_corr*: A module to calculate the correlation coefficient between two RDMs, based on different algorithms, including Pearson correlation, Spearman correlation, Kendalls tau correlation, cosine similarity, and Euclidean distance.
*corr_cal_by_rdm*: A module to calculate the representational similarities among the RDMs under different modes.
*corr_cal*: A module to conduct one-step RSA between two different modes data.
*corr_to_nii*: A module to save the representational analysis results in a .nii file for fMRI.
*stats_cal*: A module to calculate the statistical results.
*rsa_plot*: A module to plot the results from representational analysis. It contains the functions of plotting the RDM, plotting the graphs or hotmaps with results from representational analysis by time sequence based on EEG or EEG-like (such as MEG) data, plotting the results of fMRI representational analysis (montage images and surface images).

### Calculate the RDMs

An RDM is a typical approach for comparing representations in neural data. By extracting data from two different conditions and calculating the correlations between them, we will obtain the similarity between the two representations under the two conditions. Subtract the obtained similarity index from 1 and get the values of the dissimilarity index in RDM (**Figure 2**). In Fig 2, Different grating stimuli were observed to product different neural activity signals, and the value in RDM presented the dissimilarity of neural activities between two different stimuli. As shown in the figure, the closer the two grating orientations were, the lower the dissimilarity would be. In a typical object recognition experiment, humans and monkeys were allowed to watch the same sets of visual stimuli (Kriegeskorte, 2008). Researchers calculated the humans’ RDM based on fMRI data and the monkeys’ RDM based on electrophysiological data. The results indicated that the neural patterns in RDMs were similar when humans and monkeys observed stimuli belonged to the same category (animate or inanimate).

**Figure 2.**
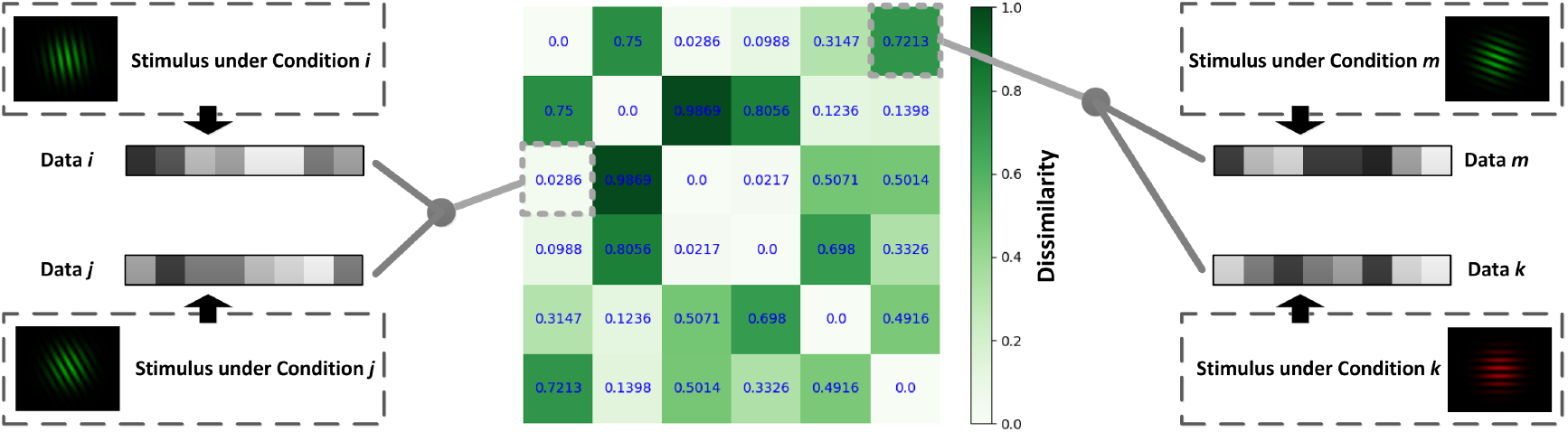
Schematic diagram for calculating the RDM. Different data can be obtained under different conditions. The value of the point [*i*, *j*] in RDM is obtained by calculating the dissimilarity (1-correlation coefficient *r*) between the data under condition *i* and that under condition *j*.

The application of calculating RDM is not restricted. NeuroRA achieves computations based on multiple modal neural data from behavioral data to brain imaging data (**Figure 3**).

**Figure 3.**
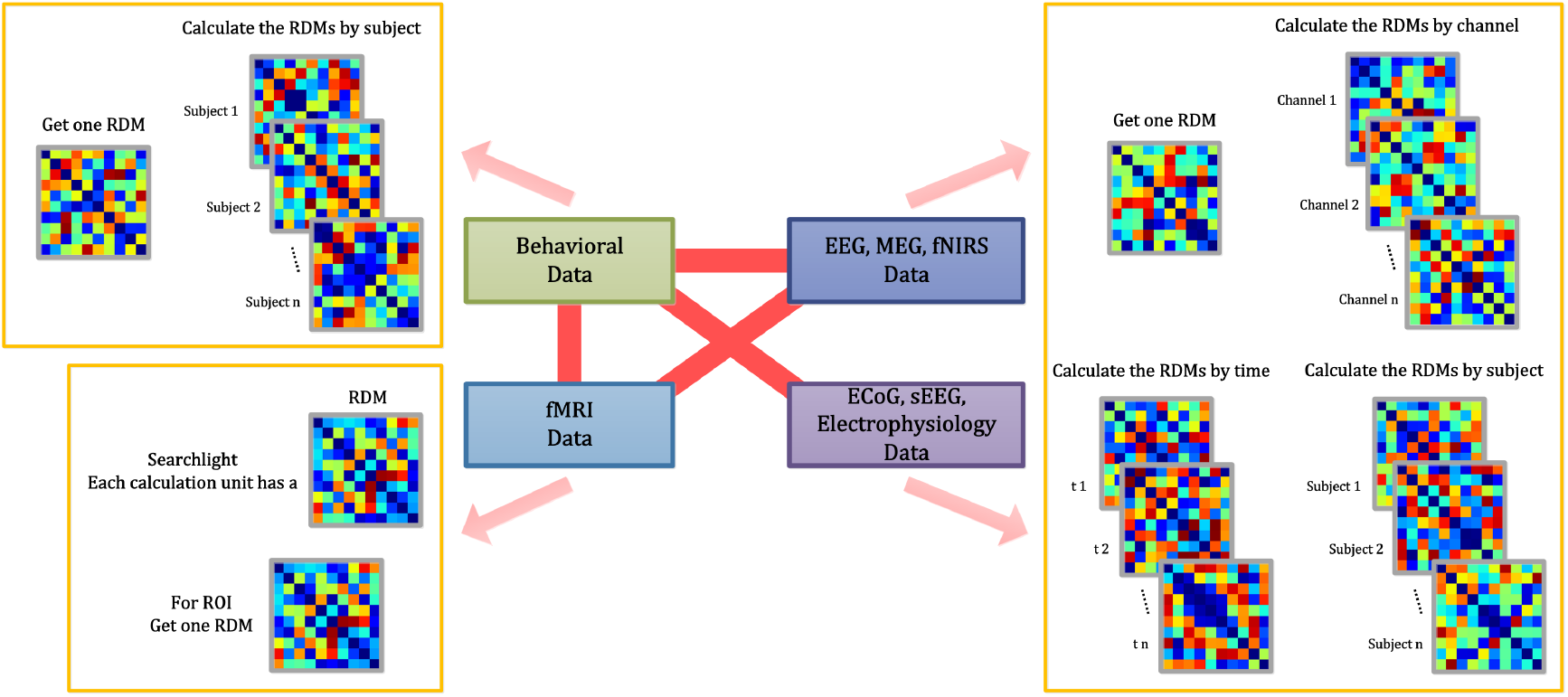
Range of the feature of calculating the RDM. NeuroRA is capable of calculating data in a variety of modes, including behavior, fMRI, EEG, MEG, fNIRS, ECoG, sEEG, and electrophysiology. The red bold lines denote the ability to perform calculations between two modes. The example referred to by the pink clip denotes the alternative calculation methods of the corresponding mode.

Assuming that the measured data from a certain condition under a total of *n* experimental conditions are denoted as *d*_1_, *d*_2_, …, *d*_*n*_, then the following RDM of *n*×*n* can be calculated by the corresponding function under the rdm_cal module from our toolkit:

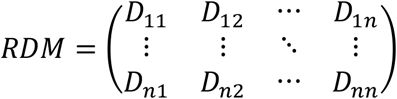

where *D*_*ij*_ denotes the dissimilarity between the data under condition *i* and that under condition *j*. The dissimilarity (*D*_*ij*_) is calculated as follows:

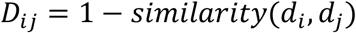

In certain cases, researchers must calculate RDM separately for each participant, or they may calculate RDM independently for each channel or each time point (Henriksson et al., 2019; Hall-McMaster et al., 2019). We resolve these issues at the beginning of NeuroRA. The toolkit provides several input parameters in the functions that allow users to make the corresponding changes in order to get RDM/RDMs by different subjects or channels or time-points or searchlight (for fMRI) or specific ROI (for fMRI) (**Figure 3**). Users can change the calculation parameters according to their requirements, and consequently, the corresponding output formats change.

### Representational analysis among different RDMs

NeuroRA provides a convenient way to calculate cross-modality similarity by computing the similarities between two RDMs from different modalities. We offer several solutions to calculate the similarity (or correlation coefficient), including Pearson correlation, Spearman correlation, Kendalls tau correlation, cosine similarity, and Euclidean distance. Users can freely change parameters to select different computing methods.

In these calculation process, we first reform the square matrices into vectors and then calculate similarities (**Figure 4**). Some previous studies calculated the correlation coefficient between two RDMs by using the diagonal values, which makes the result deceptively high (Ritchie et al., 2017). We avoid this by removing the diagonal values and include only half of the matrix to reduce the duplication, as the upper and lower halves of the RDM are actually identical (**Figure 4**).

**Figure 4.**
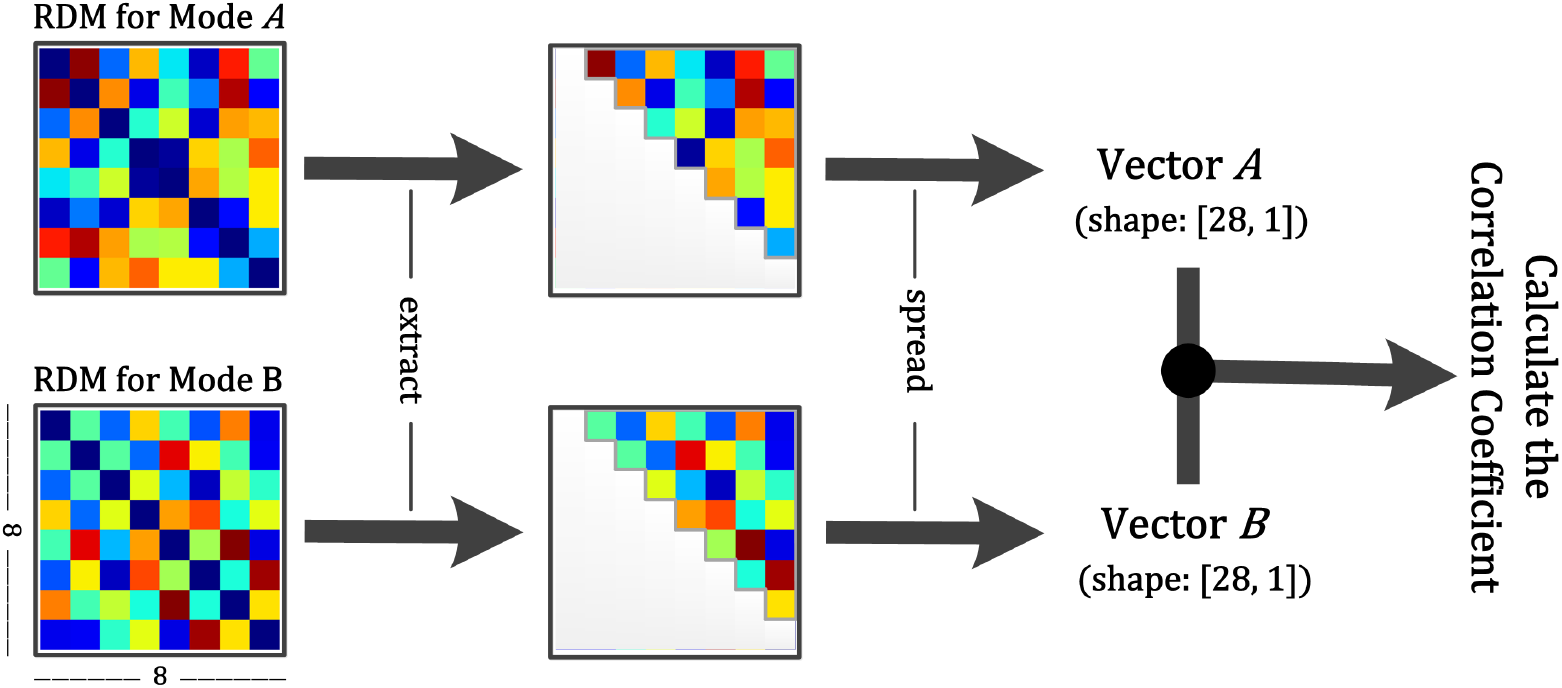
Schematic diagram for calculation between two RDMs. Step 1: Obtain two RDMs from different modal data. Step 2: Extract the points of the upper diagonal (within the grey line). Step 3: Spread them as vectors. Step 4: Calculate the correlation coefficient or conduct a permutation test between two vectors. The former returns the correlation coefficient and the *p*-value, and the latter returns only the *p*-value.

Furthermore, NeuroRA provides a permutation test to determine whether the two RDMs are related. The permutation test is based on random shuffling of data and is suitable for data with a small sample size (Efron and Tibshirani, 1994). We first shuffle the values in the two RDMs and re-calculate the similarity matrix between the two RDMs. By repeating this procedure 5000 times (the number of iterations here can be defined by users), we get the final *p*-values from this permutation distribution.

NeuroRA can also perform calculations based on multiple RDMs. Consequently, we can expand it to conditions with multiple RDMs. When you obtain a behavioral RDM from behavioral data and wish to compare it with other modal data, a problem may arise as more than one RDM can be obtained based on other modal data, such as EEG or fMRI. Our toolbox provides “searchlight” computation to perform correlation analysis between RDM from one Mode (behavior, or any of neural data) and RDM from other Modes (each individual brain region, time interval, or others) one by one (**Figure 3**). For example, calculations based on EEG data can obtain one RDM per channel or time interval or both (**Figure 5**). **Table 2** is a script example for using NeuroRA to calculate the similarities between behavioral RDM and EEG RDM per channel. Another simple example is when the users want to see which brain regions are highly correlated with behavioral performance or a coding model, they can get one behavioral/model RDM based on behavioral response time or accuracy and they may also get many fMRI RDMs from different regions. Users can put these two kinds of RDMs (behavioral/model RDM and fMRI RDMs) into our function, and they will get results showing the regions that are highly correlated with behavioral/model patterns based on thresholds of significance (p value) or correlation values set by users. (**Tabel 3**, more details on fMRI calculation are described in the next section). These convenient functions of ergodic computation cover the vast majority of cross-modal research needs.

**Table 2.** Scripts of representational analysis between behavioral data and EEG data for each channel in NeuroRA. Users can input data from different modes and obtain the correlation between results of the two modes. If users want to conduct calculation for each time-points, they can set the parameters: *time_opt*, *time_win* & *time_step* in function *eegRDM* and *bhvANDeeg_corr*.

|  |  |  |
| --- | --- | --- |
| Scheme 1 | 1 | `from neurora.rdm_cal import bhvRDM, eegRDM` |
|  | 2 | `from neurora.corr_cal_by_rdm import rdms_corr` |
|  | 3 |  |
|  | 4 | `# calculate the behavioral RDM for each subject` |
|  | 5 | `# the shape of bhv_data should be [n_conditions, n_subjects, n_trials]` |
|  | 6 | `# the shape of bhv_rdms will be [n_subjects, n_conditions, n_conditions]` |
|  | 7 | `■ bhv_rdms = bhvRDM(bhv_data, sub_opt=1)` |
|  | 8 | `# calculate the eeg RDMs for each channel & each subject` |
|  | 9 | `# the shape of eeg_data should be [n_conditions, n_subjects, n_trials, n_channels, n_times]` |
|  | 10 | `# the shape of eeg_rdms will be [n_subjects, n_channels, n_conditions, n_conditions]` |
|  | 11 | `■ eeg_rdms = eegRDM(eeg_data, sub_opt=1, chl_opt=1)` |
|  | 12 | `# initialize the correlation coefficients` |
|  | 13 | `■ corrs = np.zeros([n_subjects, n_channels, 2], dtype=np.float)` |
|  | 14 | `# calculate the correlation coefficients between behavioral RDM and eeg RDMs` |
|  | 15 | `# the shape of corrs is [n_subjects, n_channels, 2], 2 represents a r-value & a p-value` |
|  | 16 | `■ for sub in range(n_subjects):` |
|  | 17 | `■ corrs[sub] = rdm_corr(bhv_rdms[sub], eeg_rdms[sub])` |
| Scheme 2 | 18 | `from neurora.corr_cal import bhvANDeeg_corr` |
|  | 19 |  |
|  | 20 | `# calculate the correlation coefficients between behavioral RDM and eeg RDMs` |
|  | 21 | `■ corrs = bhvANDeeg_corr(bhv_data, eeg_data, sub_opt=1, chl_opt=1)` |

**Figure 5.**
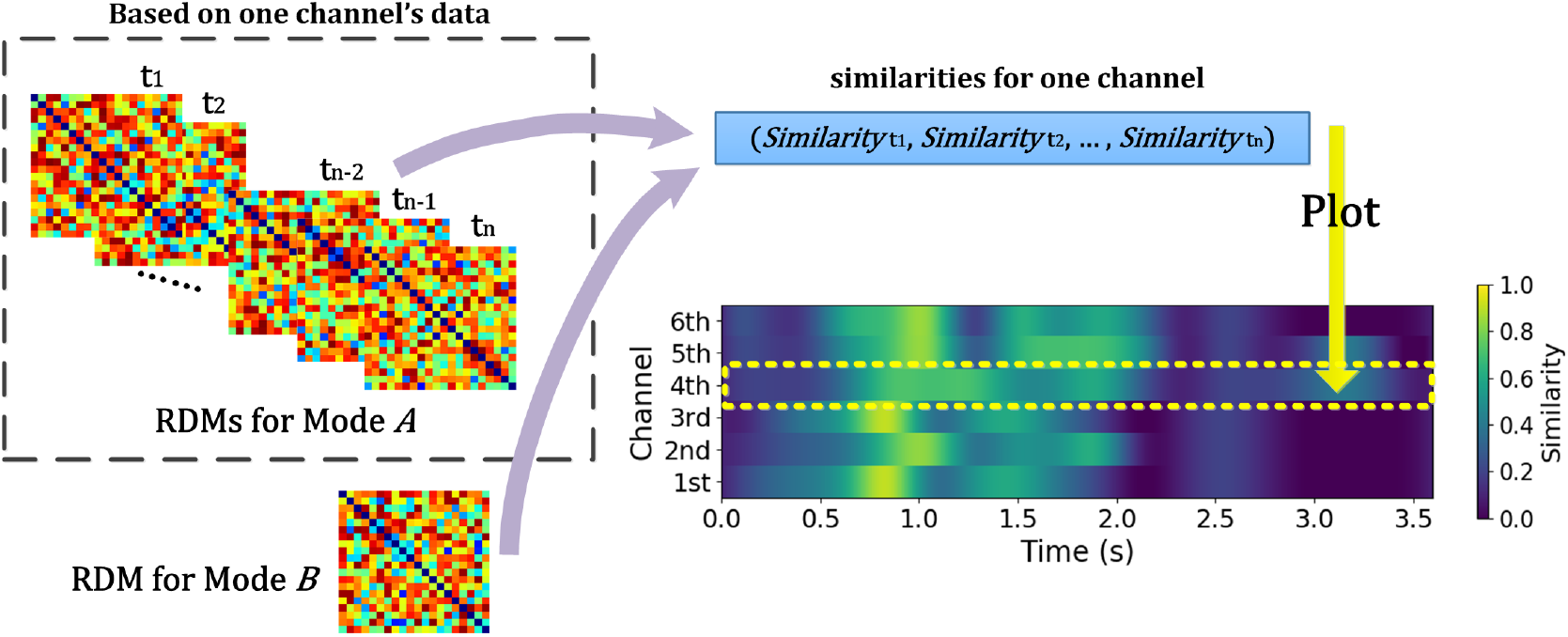
Schematic diagram for calculating similarities between RDM from different Modes across time and channel for EEG and EEG-like (such as MEG, sEEG, et al.) data. NeuroRA calculates the similarities between RDMs for mode A (EEG and EEG-like data) and one RDM for mode B (such as behavior). Such calculation can be performed across each time-point and each channel. Each value in time-channel result-image (bottom right) corresponds to a similarity index (for example, the Pearson correlation) between RDMs from two Modes.

To maximally simplify users’ experience, our toolbox offers a one-step option between different modes (**Tabel 2 Scheme 2** is a one-step example for calculating a similarity index between behavior and EEG). Users can input data from two modalities, and the toolbox will return the final results of representation analysis. This will be very convenient and efficient when the users do not need to obtain the RDMs in the intermediate stages.

### Searchlight across the whole brain and save results as a NIfTI file for fMRI data

In the field of cognitive neuroscience, fMRI research constitutes a large proportion. Researchers typically wish to calculate RDMs for different brain regions. Users can conduct representational analysis for ROIs or searchlight across the whole brain for fMRI data based on NeuroRA (**Figure 6**).

**Figure 6.**
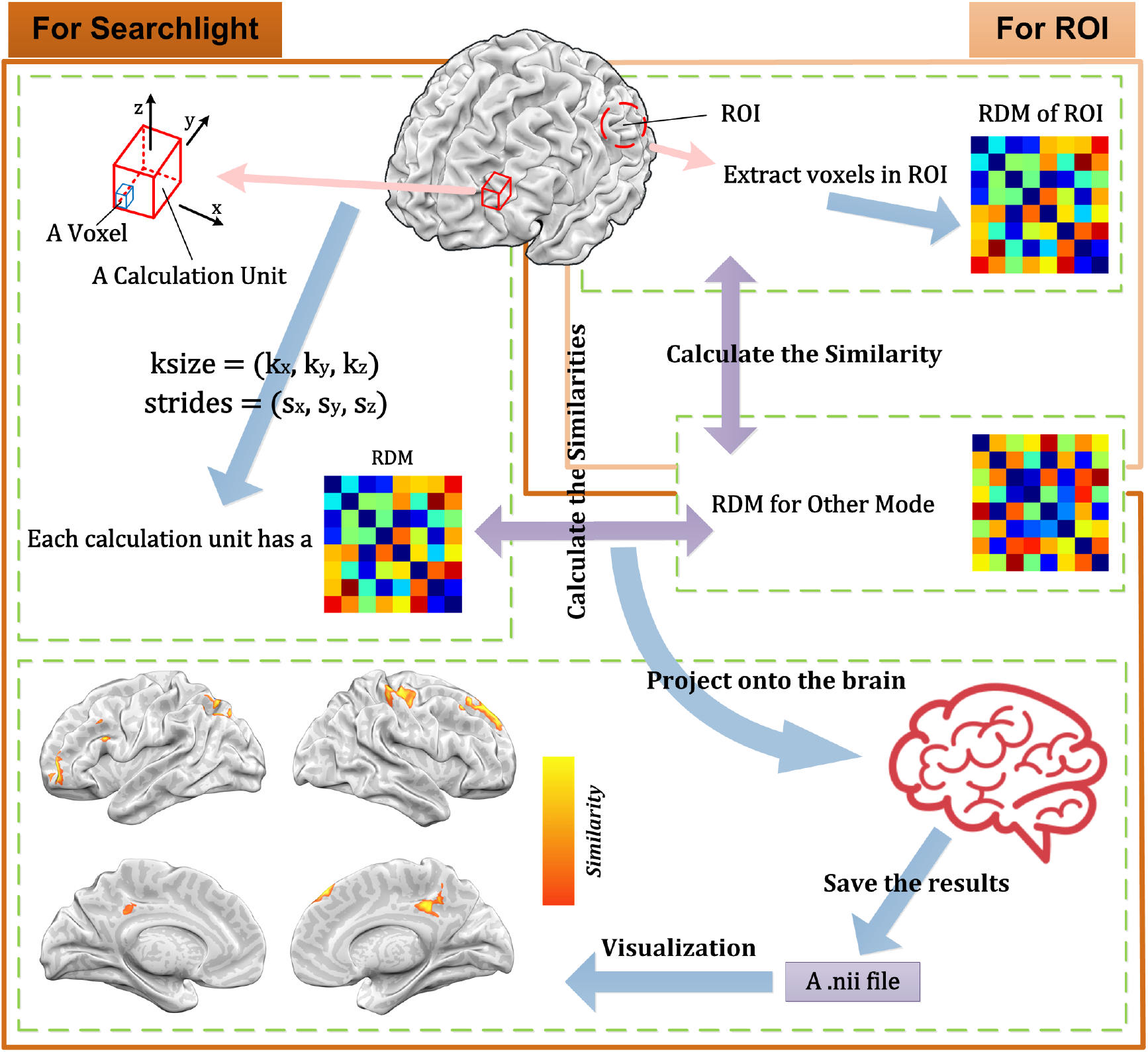
Schematic diagram for searchlight calculation for fMRI data based on NeuroRA. Inside the area with dark brown borders is the calculating process for each searchlight process. Inside the area with light brown borders is the calculating process for each ROI. For each searchlight step, users define the size and strides of the calculation unit. After computations between the RDMs within the searchlight blob for fMRI and the RDM for other modes (e.g. behavioral data, computer simulated data), a NIfTI file can be obtained. At the bottom left is a demo of the resulting NIfTI file drawn with NeuroELF (http://neuroelf.net), and color-coded regions indicate strength of representation similarity between two modes. For each ROI, users can calculate the RDM based on the voxels in ROI and get similarity between ROI RDM and the RDM for other modes.

For each searchlight step, users can customize the size of the basic calculation unit [*k*_*x*_, *k*_*y*_, *k*_*z*_]. Each *k* indicates the number of voxels along the corresponding axis. The strides between different calculation unit must be decided as [*s*_*x*_, *s*_*y*_, *s*_*z*_]. The strides refer to how far the calculation unit is moved before another computation is made. Each *s* indicates how many voxels exist between two adjacent calculation units along the corresponding axis. For the fMRI data of size [*X*, *Y*, *Z*], the kernel size is usually set to be more than one voxel, so each voxel can exist in multiple kernels (calculation units). Therefore, *N* computations are required here:

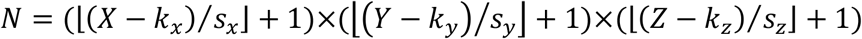

This implies that *N* RDMs must be calculated, which are each related to the corresponding calculation unit. After obtaining searchlight RDMs, users can calculate the similarities between fMRI and other modes. In NeuroRA, the final correlation coefficient of one voxel is the mean value of the correlation coefficients calculated by all kernels that contain this voxel.

**Tabel 3** is a script demo to understand how to conduct a searchlight based analysis for fMRI data. We could first calculate the fMRI RDMs within each searchlight blob, and then obtain similarities between fMRI RDMs and a coding model RDM all over the whole brain.

**Table 3.**
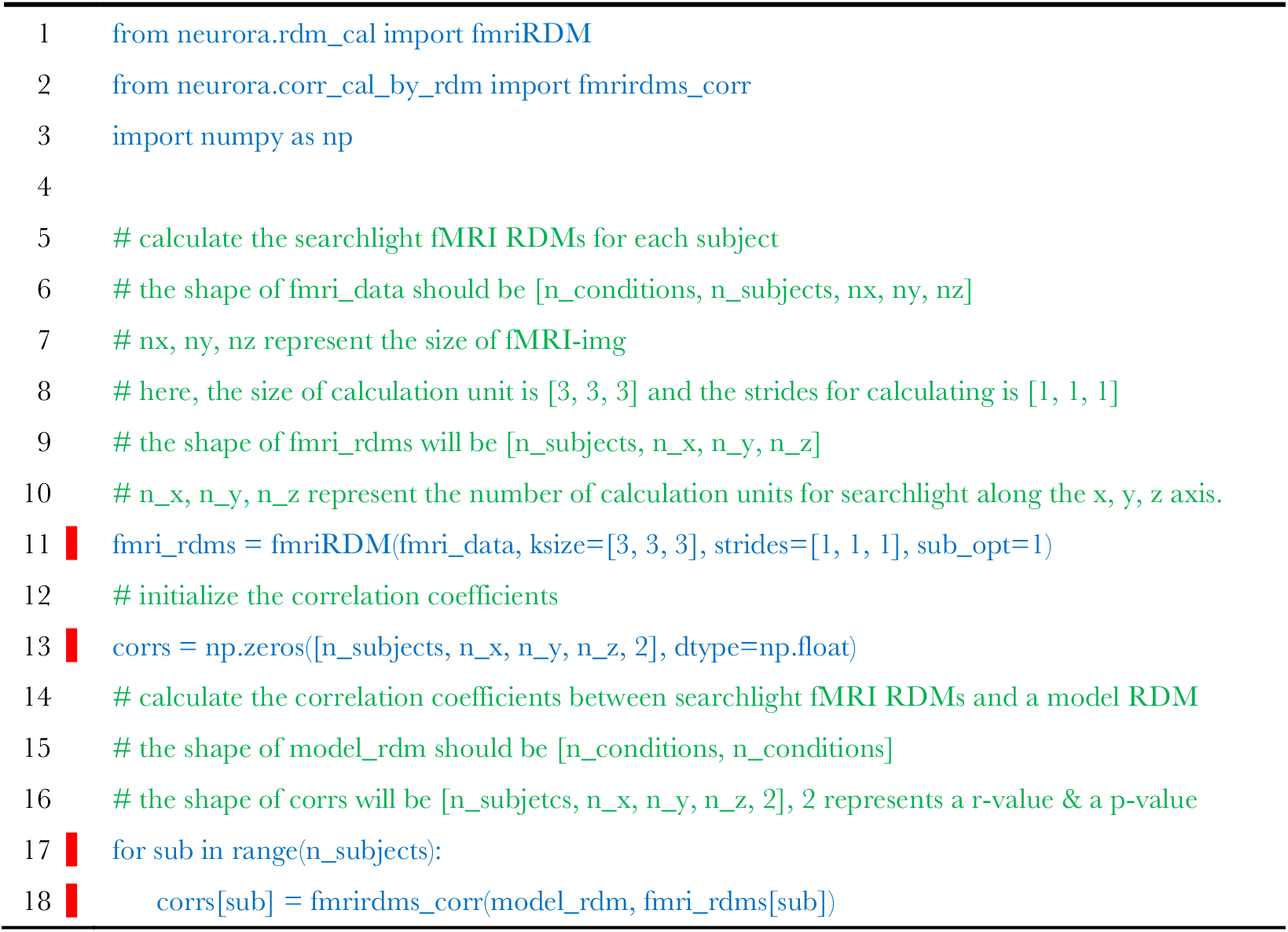
Script of searchlight representational analysis between fMRI data and a coding model in NeuroRA. The calculation parameters of fMRI data are *ksize=[3, 3, 3]* and *strides=[1, 1, 1]*. Users can just input different data and obtain the correlation results between two modes.

NeuroRA provides functions to save the results as a NIfTI file with thresholding parameters as well (**Table 4**). Users can set certain thresholds for *p*-values, *r*-values or *t*-values. Also, users can select Family-Wise-Error (FWE) or False-Discovery-Rate (FDR) correction methods to control for multiple comparisons across the whole brain. Furthermore, users can choose whether to smooth the results, whether to plot automatically, etc. For example, if the threshold for *p* value is set as 0.05, the final NIfTI file returned will be filtered with *p* < 0.05, and all voxels with *p*>=0.05 will be set as 0.

**Table 4.**
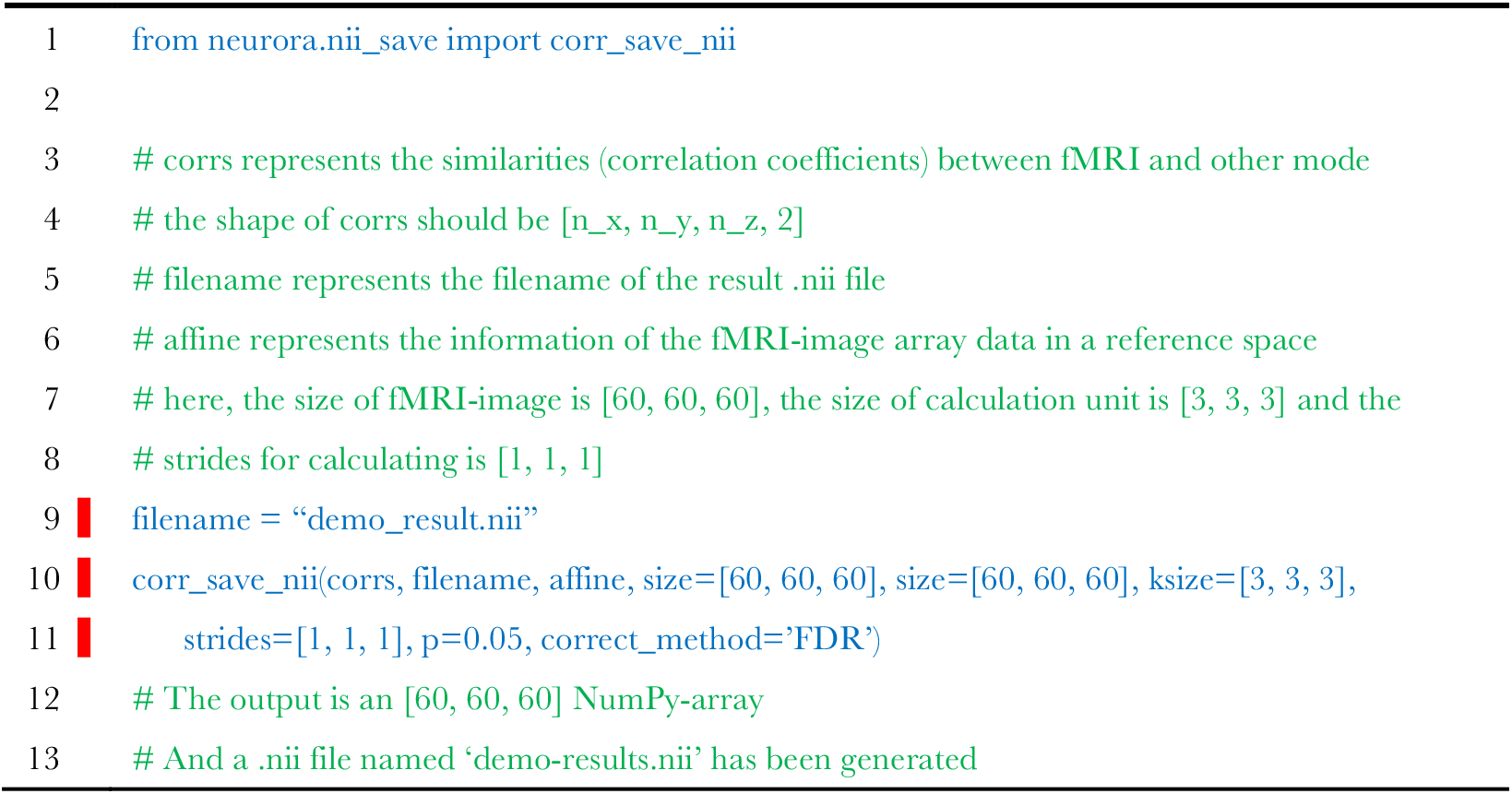
Script of saving the calculation results as a NIfTI file for fMRI data. Users can get the correlation results based on the script in **Tabel 3**. The NIfTI file can be obtained by entering some necessary parameters.

### Other Representational Analysis

In addition to RSA, users can conduct analysis of NPS, STPS and ISC with NeuroRA. Our toolkits have separate modules to conduct these calculations. Just like RSA from multiple modalities, the calculations for representational analysis above support EEG-like data as well as fMRI data. Users can calculate the results for each channel/region, each time-point from a time series, each ROI or searchlight blobs (for fMRI) as they wish by selecting different functions and setting specific parameters. These calculations are used in a similar way to calculate RDM or RSA in the above sections. Users can use *help()* (a built-in function in Python) to see and understand the detailed description of the purpose of the specific function or module, and users can get more information from our tutorial documents.

### Statistical Analysis

NeuroRA provides functions for statistical analysis based on the calculations mentioned above. The inputs are the r-value & p-value maps for each subject, and the output will be the statistical results (a t-value & p-value map) (Table 5). Besides, we add permutation test to all processes of statistical analysis. This means the statistical significance could be assessed through a permutation test by randomly shuffling the data and calculated the results for many iterations (for example 5000) to draw a distribution. Real data exceeding 95% of the distribution are regarded significant. Users can independently choose to use permutation test or not and change the iteration number by set parameters in related functions.

**Table 5.** Example of statistical analysis for channel-time based EEG calculation and searchlight fMRI calculation.

| Type of Calculation | Example Script |
| --- | --- |
| *channel-time based EEG calculation* | `from neurora.stats_cal import stats # the shape of corrs should be [n_subs, n_chls, n_ts, 2] stats(corrs, permutation=True, iter=1000) # The output is an [n_chls, n_ts, 2] NumPy-array # 2 represents a t-value and a p-value` |
| *searchlight fMRI calculation* | `from neurora.stats_cal import stats_fmri # the shape of corrs should be [n_subs, n_x, n_y, n_z, 2] stats_fmri(corrs, permutation=True, iter=10000) # The output is an [n_x, n_y, n_z, 2] NumPy-array` |

### Visualization of Results

NeuroRA provides several functions to visualize the results in *rsa_plot* module (**Figure 7**). These allow users to use our toolkit to plot whatever they select.

**Figure 7.**
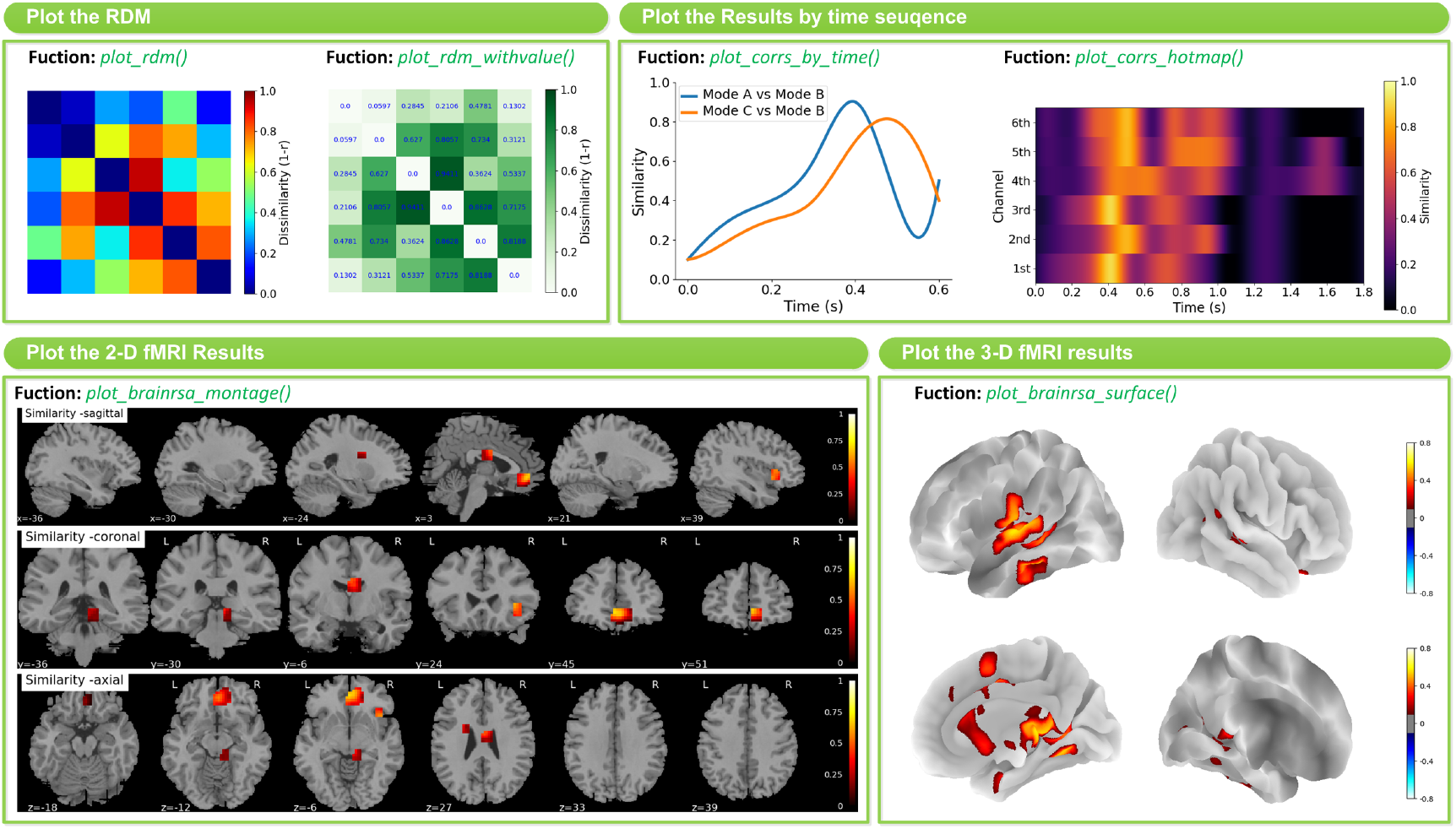
Typical features of visualization. Left-top: Plot the RDM by function *plot_rdm()* and *plot_rdm_withvalue()*. Right-top: Plot the results by time sequence by function *plot_corrs_by_time()* and *plot_corrs_by_hotmap()*. Left-down: Plot the 2-D fMRI results by function *plot_brainrsa_montage()*. Right-down: Plot the 3-D fMRI results by function *plot_brainrsa_surface()*.

The basic option is to visualize RDMs by function *plot_rdm()* or *plot_rdm_withvalue()*. Users can select different colormaps and set up the threshold for correlation coefficients to appear. The more advanced option for EEG and EEG-like data is to visualize the results across different time points by function *plot_corrs_by_time()* and *plot_corrs_hotmap()*, etc. The similarity index is obtained from each time point and cascaded together. The former is a graph and the latter is a hotmap. Also, NeuroRA has options for plotting fMRI results on a brain. Users can use *plot_brainrsa_montage()* and some other functions to plot 2-D results and use *plot_brainrsa_surface()* to plot 3-D results. The implementation of visualization requires the Pyplot module in the Matplotlib and nilearn package.

We also provide several code demos in NeuroRA on the publicly available datasets. One is based on visual-92-categories-task MEG datasets (Cichy et al., 2014). The other one is based on Haxby fMRI datasets (Haxby, 2001). In these demos, user can learn how to use NeuroRA to perform representational analysis and plot the essential results, including calculating RDMs from different time points (**Fig 8a**), correlations over the time series (**Fig 8b**), ROI calculation (**Fig 8c**, here use a ventral temporal mask), searchlight calculation between the brain activities and a ‘animate-inanimate’ coding model (**Fig 8d**) and so on. These demos contain several critical sections: loading data & preprocessing, calculating RDMs, calculating the neural similarities or similarity matrix and plotting. Users can download the tutorial in NeuroRA website and find further details.

**Figure 8.**
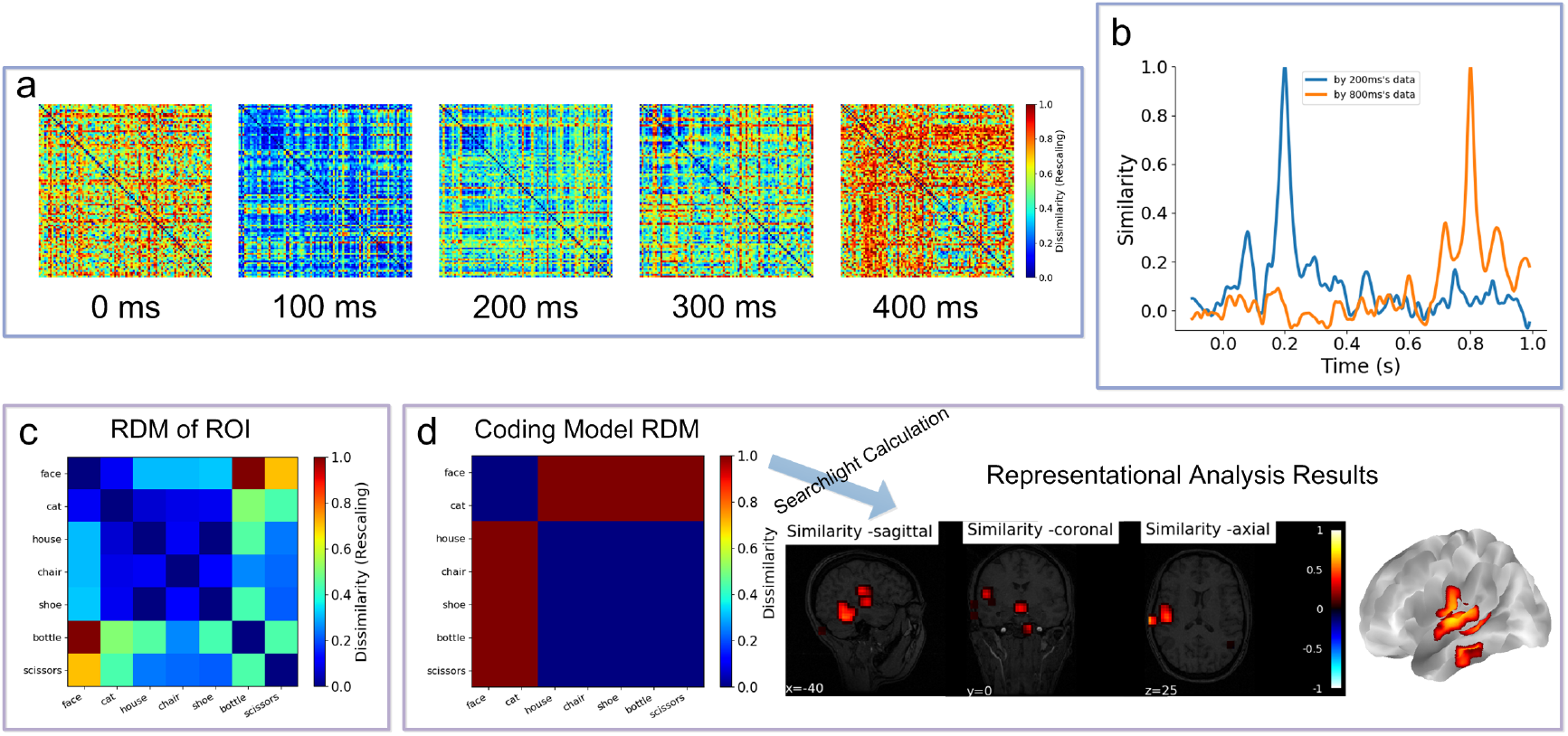
Demo results. (a) The RDMs of 0ms, 100ms, 200ms, 300ms and 400ms based on all 302 channels’ MEG data. (b) Use the neural representations of 200ms and 800ms to calculate the similarities with all time-points’ neural representations. (c) The RDM for ROI (ventral temporal). (d) The searchlight results between a ‘animate-inanimate’ coding model RDM and searchlight RDMs. In this coding model RDM, we assume that there are consistent representations among animate objects and among inanimate objects.

## User Support

To report any bugs in the code or submit any queries or suggestions about our toolbox, users can use the issue tracker on GitHub: https://github.com/neurora/NeuroRA/issues. We will reply and act accordingly as soon as possible.

## Discussion

RSA provides a novel way of observing big data, which is powerful in the field of cognitive neuroscience. An increasing number of studies have used such multivariate analysis to obtain novel information that could not be observed through univariate analysis (Mahmoudi et al., 2012; Sui et al., 2012; Haxby et al., 2014). More importantly, research based on different modalities must be assessed simultaneously, and RSA offers a perfect way to quantitatively compare results from different modalities with distinctive dimensions, resolutions, and even different species (Cichy and Pantazis, 2017; Salmela et al., 2016).

In the present study, we have developed a Python-based toolbox that can perform representation analysis for neural data from many different modalities. Compared with other toolkits or modules that can also implement RSA, our toolbox has much wider application, stronger computational performance, and more convenient and complete functions, especially for the analysis of multi-modal data and cross-modal comparisons. Moreover, it is open source, free to use, and cross-platform.

For detailed information on each module and function in our toolbox, including the format of input data, the choice of parameters, and the format of output data, users can refer to our toolbox tutorial. To further understand the specific implementation of each function in the toolbox, people can read the source code directly. If users encounter any problems or difficulties during use, they can consult NeuroRA’s tutorials and email our developers.

Although NeuroRA has already realized the essential functions for RSA analysis, there are still several limitations to be addressed in the future. First, NeuroRA is not yet designed to process the raw data. However, users can utilize other toolbox such as EEGLAB (Delorme and Makeig, 2004), MNE (Gramfort et al., 2013), and Nibabel (Brett et al., 2016), to import data and transfer them into a format fit for NeuroRA. We plan to develop an integrated format conversion function in the next version. Second, there is still significant room for improving the computational performance of NeuroRA, especially in the iterative calculation of fMRI data, which is often slow. This is due to nested loops in the code structure when we need to traverse the data from the entire brain and iterate the random shuffle many times. In the future, we will reduce the time by optimizing functions with GPUs. Third, there is currently no graphical user interface (GUI) right now, which we plan to design and implement based on PyQt in a future version. Users could then obtain the final results by dragging the data file to a specific location in the GUI with the mouse and filling in the relevant parameters.

Through NeuroRA, for the first time, we provide a method for researchers to perform representation analysis with neural data from many different modalities. However, this is only a starting point. With the development of the algorithm to compute representational analysis, we will include new methods, as well as new visualization tools. We hope that many interesting findings can be observed through our toolbox and we would like to collaborate with researchers that are interested in this tool to improve the toolbox further.

## Information Sharing Statement

NeuroRA is available at https://pypi.org/project/neurora/. The website for NeuroRA is https://neurora.github.io/NeuroRA/, and the tutorial of the toolbox can be download here. The code for our toolbox NeuroRA can be accessed on GitHub: https://github.com/neurora/NeuroRA.

